# Epigenetic factors coordinate intestinal development

**DOI:** 10.1101/399410

**Authors:** Julia Ganz, Ellie Melancon, Catherine Wilson, Angel Amores, Peter Batzel, Marie Strader, Ingo Braasch, Parham Diba, Julie A. Kuhlman, John H. Postlethwait, Judith S. Eisen

## Abstract

Intestinal epithelium development depends on epigenetic modifications, but whether that is also the case for other intestinal tract cell types remains unclear. We found that functional loss of a DNA methylation machinery component, *ubiquitin-like protein containing PHD and RING finger domains 1 (uhrf1),* leads to reduced enteric neuron number, changes in neuronal morphology, and severe intestinal smooth muscle disruption. Genetic chimeras revealed that Uhrf1 functions both cell-autonomously in enteric neuron progenitors and cell-non-autonomously in surrounding intestinal cells. Uhrf1 recruits the DNA methyltransferase Dnmt1 to unmethylated DNA during replication. Dnmt1 is also expressed in enteric neuron and smooth muscle progenitors. *dnmt1* mutants show a strong reduction in enteric neuron number and disrupted intestinal smooth muscle. Because *dnmt1*;*uhrf1* double mutants have a similar phenotype to *dnmt1* and *uhrf1* single mutants, Dnmt1 and Uhrf1 must function together during enteric neuron and intestinal muscle development. This work shows that genes controlling epigenetic modifications are important in coordinating intestinal tract development, provides the first demonstration that these genes are important in ENS development, and advances *uhrf1* and *dnmt1* as potential new Hirschsprung disease candidates.

**Summary:** This work provides evidence that DNA methylation factors are important in all cell types that contribute to development of a functional intestine.

## Introduction

Proper organ function requires coordinated development of multiple cell types over an appropriate temporal window. For example, development of a functioning intestinal tract involves coordination among intestinal epithelial cells derived from endoderm, muscle cells derived from mesoderm, and enteric neurons and glia derived from ectoderm. Cells from each of the three germ layers initially migrate and then proliferate, prior to their differentiation, which occurs essentially contemporaneously (Ganz, 2018; Ganz et al., 2016; Hao et al., 2016; Wallace et al., 2005a). Endoderm and mesoderm progenitor cells initially co-mingle (Gays et al., 2017; Warga and Nusslein-Volhard, 1999; Zorn and Wells, 2009), whereas the ectoderm that generates the neural crest cells that give rise to the enteric nervous system (ENS) is positioned far from the nascent endoderm and mesoderm. Thus, these ENS precursors must migrate for a significant distance to reach the developing intestinal tract before they can begin to differentiate and innervate it (Ganz, 2018; Lake and Heuckeroth, 2013). Signaling among these distinct cell types, or their progenitor cells, is critical for coordinated intestinal development. The intestinal epithelium influences development and differentiation of intestinal smooth muscle precursors (ISMPs) and intestinal smooth muscle cells (ISMCs), as well as development and differentiation of enteric precursor cells (EPCs) and the ENS (Fu et al., 2004; Pietsch et al., 2006; Reichenbach et al., 2008; Sukegawa et al., 2000). The lateral plate mesoderm (LPM) from which ISMPs and ISMCs arise influences intestinal epithelial development and EPC migration and differentiation (Fu et al., 2004; Graham et al., 2017; Hao et al., 2016; Mwizerwa et al., 2011; Natarajan et al., 2002; Puzan et al., 2018; Reichenbach et al., 2008; Sukegawa et al., 2000). Similarly, the ENS regulates intestinal epithelial development and integrity (Neunlist et al., 2007; Neunlist et al., 2013; Puzan et al., 2018). Thus, there is interdependence of intestinal cell type development through signaling between progenitor cells of intestinal epithelium, smooth muscle, and ENS. Mutations in genes involved in development of any of these cell types can have profound and often deleterious consequences (Brosens et al., 2016; Goldstein et al., 2016; Wallace et al., 2005b; Yamamoto and Oda, 2015). For example, mutations in genes that regulate ENS development can result in disorders, such as Hirschsprung Disease (HSCR), in which the distal intestine is uninnervated and dysmotile (Heuckeroth, 2018).

A number of studies have revealed the importance of epigenetic modulation during development and functioning of the endodermal and mesodermal cell types that contribute to the intestinal tract (Elliott and Kaestner, 2015; Elliott et al., 2015; Jorgensen et al., 2018). For example, epigenetic modulation has been shown to be important for integrity of the intestinal epithelium (Marjoram et al., 2015) and proper development of intestinal smooth muscle (Jorgensen et al., 2018). Epigenetic modification via DNA methylation is a key regulator of the differential gene expression that underlies the ability of cells to develop distinct fates. DNA methylation patterns are established by the *de novo* DNA methyl transferases (Dnmt) 3a and 3b and DNA methylation patterns are maintained by Dnmt1 (Robertson and Wolffe, 2000). DNA methylation regulates intestinal epithelium formation by controlling the balance between cell proliferation and differentiation during development (Elliott and Kaestner, 2015; Marjoram et al., 2015; Sheaffer et al., 2014). Dnmt1 has also been shown to have an essential role in regulating intestinal smooth muscle differentiation, integrity, and survival (Jorgensen et al., 2018). DNA methylation has further been linked to ENS development because EPCs have decreased Dnmt expression in HSCR patients compared to controls and some HSCR patients have presumable pathogenic missense mutations in Dnmt3b (Torroglosa et al., 2014). Zebrafish mutants of *histone deacetlyase 1* (*hdac1)*, another epigenetic modifier gene, have fewer enteric neurons in addition to other neural crest defects (Ignatius et al., 2013). However, how epigenetic modifications, such as DNA methylation, affect ENS development remains poorly understood and it is also unclear how DNA methylation coordinates proper temporal development of the distinct cell types that coalesce to form the intestinal tract. Here, we investigate the role of epigenetic modulation in intestinal development by focusing on both ectodermal derivatives that form the ENS and mesodermal derivatives that form the intestinal smooth muscle.

Dnmt proteins require partners to effect DNA methylation. For example, Dnmt1 is recruited by the modular protein Ubiquitin-like protein containing PHD and RING finger domains 1 (Uhrf1) to unmethylated DNA, to maintain DNA methylation patterns after replication (Bestor, 2000; Bostick et al., 2007; Ooi and Bestor, 2008; Sharif et al., 2007). Zebrafish has been instrumental in elucidating the role of Uhrf1, as mouse mutants die early in development (Bostick et al., 2007; Muto et al., 2002; Sharif et al., 2007). Zebrafish *uhrf1* mutants show compromised intestinal barrier function resulting from disruption of the intestinal epithelium (Marjoram et al., 2015). They also show intestinal inflammation reminiscent of inflammatory bowel disease (Marjoram et al., 2015). However, little is known about how Uhrf1 influences ENS or smooth muscle cell development.

In this study, we examine the role of Uhrf1 in development of the ENS and intestinal muscle using a mutant *uhrf1* allele (*uhrf1^b1115^*, hereafter referred to as *uhrf1*^-/-^) that we isolated in a zebrafish forward genetic screen for mutants with changes in enteric neuron number (Kuhlman and Eisen, 2007). As previously reported, *uhrf1* mutants exhibit significant disruption of intestinal epithelial morphology (Marjoram et al., 2015). We demonstrate that they also exhibit severe disruption of intestinal smooth muscle, a variable reduction in enteric neuron number, and changes in enteric neuron morphology. This disruption of both EPCs and smooth muscle cells results in displacement of enteric neurons via both cell-autonomous and cell-non-autonomous mechanisms. We show that both *uhrf1* and *dnmt1* are expressed in EPCs and surrounding intestinal cell types. Consistent with the known interactions between Uhrf1 and Dnmt1, zebrafish *dnmt1* single mutants exhibit similar ENS and muscle phenotypes to *uhrf1* mutants. Our double mutant analysis demonstrated that Uhrf1 and Dnmt1 function together to regulate enteric neuron and intestinal smooth muscle development. This work provides evidence that genes controlling epigenetic modifications play an important role in coordination of intestinal development.

## Results

**Zebrafish *uhrf1* mutants have fewer enteric neurons with altered cell morphology** *uhrf1^b1115^* was originally isolated from a forward genetic screen for mutations affecting ENS development (Kuhlman and Eisen, 2007) based on a decrease in the number of ENS neurons in larvae that looked otherwise fairly normal. The mutation was identified using RAD-mapping, as described in the Methods (Fig. S1) and confirmed by non-complementation with another allele of *uhrf1* (*hi272*) (Sadler et al., 2007). There are several additional *uhrf1* mutant alleles that have been demonstrated to decrease DNA methylation (Marjoram et al., 2015; Tittle et al., 2011). To investigate the ENS phenotype in more detail, we examined the number of ENS neurons in proximal and distal intestine at 5 days post fertilization (dpf). At this stage, there are two main neuronal subtypes, those expressing neuronal nitric oxide synthase (nNOS) and those expressing serotonin (5-HT) (Uyttebroek et al., 2010). A change in the number of ENS neurons could reflect a change in either one or both of these subtypes. We counted the overall number of neurons using an antibody to Elavl which is expressed in all ENS neurons (Kuhlman and Eisen, 2007), as well as antibodies to these two neurotransmitters (Uyttebroek et al., 2010). We found that the overall number of ENS neurons was significantly reduced in both proximal and distal intestine, as a result of reductions in both subtypes (Fig. 1, 2). Notably, ENS neurons were essentially absent from the distal-most 100 μm of the intestine (Fig. 1L, Fig. 2D,H).

**Figure 1.**
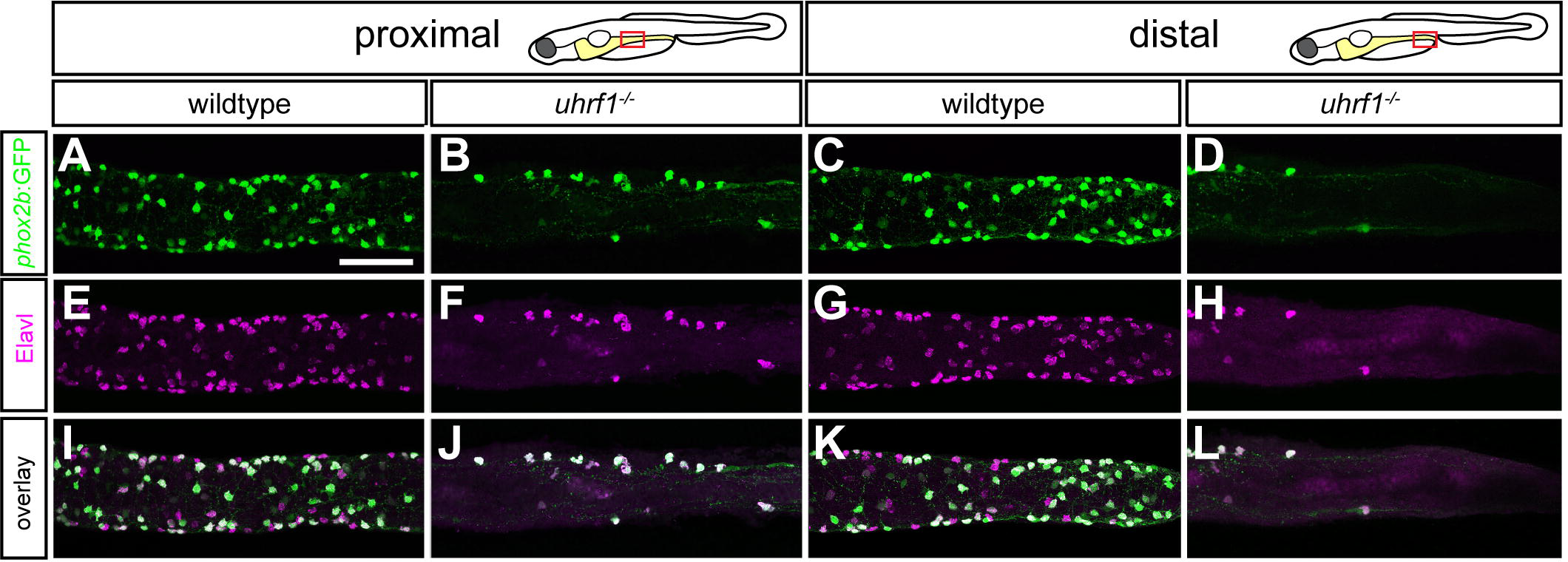
*uhrf1* mutants have fewer neurons in both proximal and distal intestine. In the proximal intestine, *uhrf1* mutants (B,F,J) have fewer enteric neurons [*phox2b:EGFP* (green), Elavl (magenta)] than wildtype siblings (A,E,I). In the distal intestine, *uhrf1* mutants (D,H,L) have fewer enteric neurons [*phox2b:EGFP* (green), Elavl (magenta)] than wildtype siblings (C,G,K). A-L: Confocal images of dissected guts at the levels indicated. Scale bar = 50μm.

**Figure 2.**
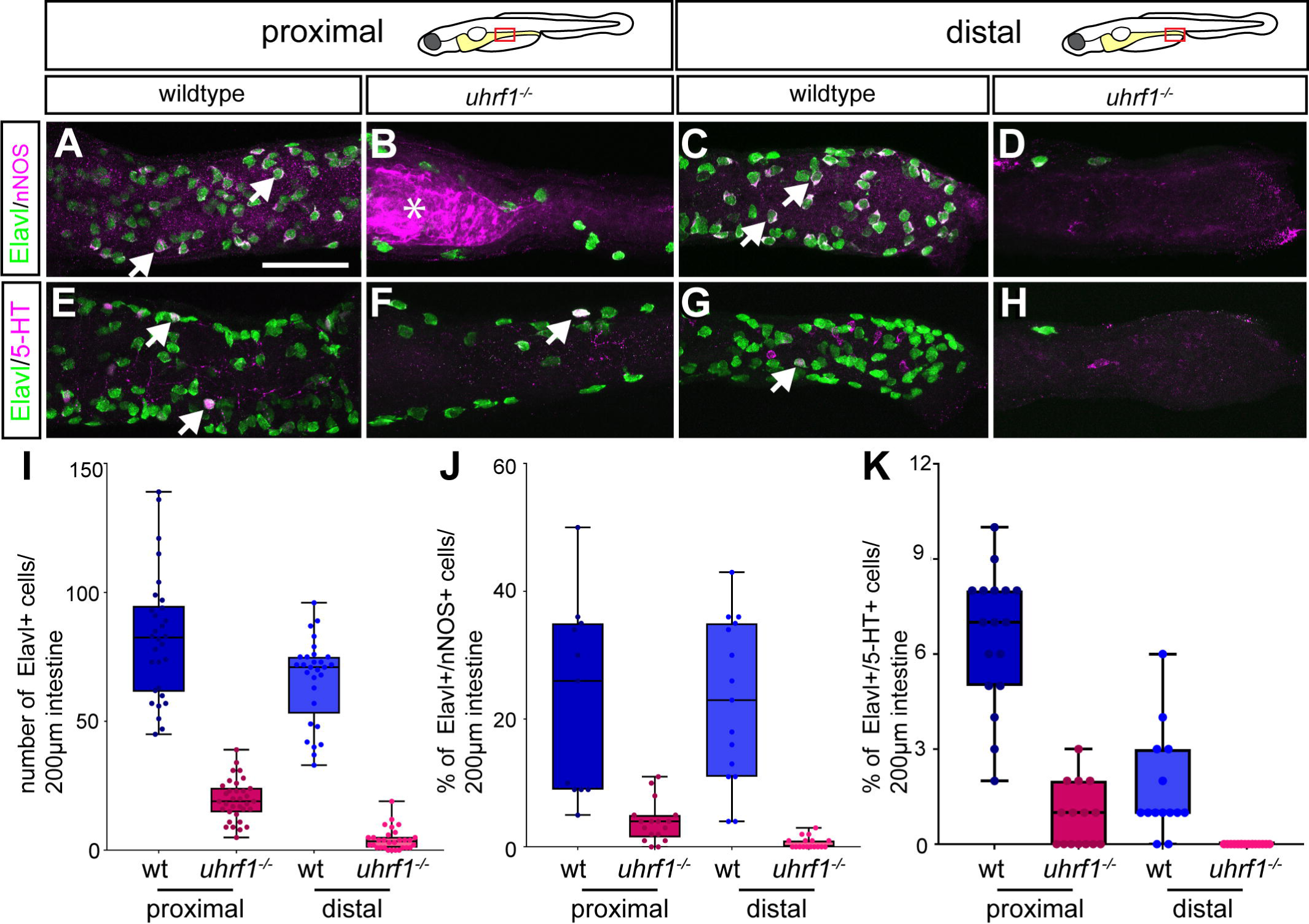
*uhrf1* mutants have fewer neurons of two neuronal subtypes both in proximal and distal intestine. In both proximal and distal intestine, *uhrf1* mutants (B,F; D,H) have fewer enteric neurons and both neuronal subtypes show similar reductions [Elavl (green), nNOS or 5-HT (magenta), arrows point to examples of nNOS or 5-HT positive neurons] compared to wildtype siblings (A,E; C,G). (I) Quantification of Elavl positive cells per 200μm of proximal or distal intestine in wildtype (blue) or mutant (red) [wildtype n=30 (proximal), n=29 (distal); *uhrf1^-/-^* n=29 (proximal), n=30 (distal)]. (J-K) Quantification of percentage of Elavl and nNOS (J) and 5-HT (K) positive cells per 200μm proximal of distal intestine in wildtype (blue) and mutant (red) [nNOS: wildtype n=11 (proximal), n=15 (distal); *uhrf1^-/-^* n=17 (proximal), n=17 (distal); 5-HT: wildtype n=17 (proximal), n=14 (distal); *uhrf1^-/-^* n=15 (proximal), n=13 (distal)]. Asterisk in B indicates autofluorescent background common to *uhrf1* mutant intestines. A-H: Confocal images of dissected intestines at the levels indicated. Scale bar = 50μm in A-H.

### Enteric progenitors and surrounding intestinal cells express *uhrf1* and *dnmt1*

As a first step toward elucidating the role of *uhrf1* in the developing intestinal tract, we examined the *uhrf1* expression pattern by *in situ* hybridization. Expression of *uhrf1* has been documented in zebrafish embryos in developing endoderm (Sadler et al., 2007). Previous studies showed that Uhrf1 is required for normal development of the intestinal epithelium in zebrafish, but did not investigate its role in development of other intestinal cell types such as intestinal smooth muscle cells or ENS neurons (Marjoram et al., 2015). EPCs, endoderm, and developing mesoderm are in such close proximity during intestinal development (Wallace et al., 2005a) that previous studies did not differentiate the expression pattern of *uhrf1* among these cell types. The transgene Tg(*phox2b*:*EGFP*) is expressed in migrating EPCs beginning around 30-32 hours post fertilization [hpf, (Shepherd et al., 2004; Taylor et al., 2016)] allowing identification of EPCs among surrounding intestinal cells. We evaluated *uhrf1* RNA expression in embryos homozygous for the *phox2b:EGFP* transgene at 48 hpf and performed immunohistochemistry to detect GFP expression in EPCs (Fig. 3A). Similar to Sadler and colleagues (2007), we observed *uhrf1* transcript in the tectum, retina, branchial arches, and developing endoderm (Fig. 3A and data not shown). We also observed *uhrf1* expression in *phox2b*:*EGFP*-positive EPCs and surrounding intestinal cells including presumptive ISMPs, suggesting that *uhrf1* plays a role in both ENS and intestinal smooth muscle development (Fig. 3A), as well as in intestinal epithelium development, as previously shown (Marjoram et al., 2015).

**Figure 3.**
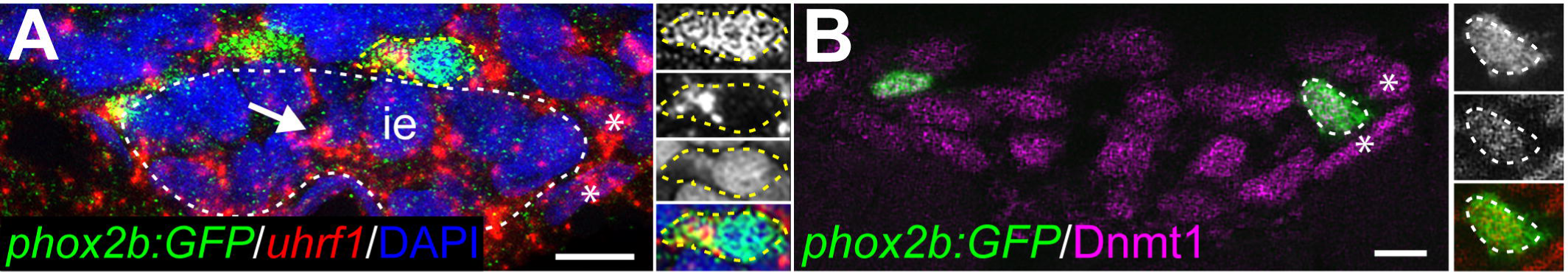
*uhrf1* and Dnmt1 are expressed in enteric, epithelial, and smooth muscle progenitors during development. (A) At 48 hpf, *uhrf1* (red) is expressed in *phox2b:EGFP* (green) positive EPCs and also in DAPI-positive ISMPs (red, blue, asterisks) and epithelial progenitors (red, blue, arrow). White dashed line indicates intestinal epithelium (ie). Insets show close-ups of outlined cell (yellow dashed line), GFP, *uhrf1*, DAPI, and overlay from top to bottom. (B) At 34 hpf, Dnmt1 (magenta) is expressed in *phox2b:EGFP* (green) positive EPCs and ISMPs (asterisks). Insets show close-ups of outlined cell, GFP, Dnmt1, and overlay from top to bottom. A,B: cross-sections of embryos. Scale bar = 10μm.

Uhrf1 is necessary to recruit Dnmt1 to unmethylated DNA during replication (Bostick et al., 2007; Muto et al., 2002; Sharif et al., 2007), and Dnmt1 is necessary to maintain intestinal epithelial progenitor cells (Elliott et al., 2015). Given this interaction, and that expression of Dnmt1 has been documented in zebrafish embryos in developing endoderm (Liu et al., 2015; Rai et al., 2006), we wondered whether Dnmt1 protein is also expressed in EPCs and surrounding intestinal cell types. To address this question, we immunostained transverse sections of *phox2b:EGFP*-expressing embryos at 34 and 36 hpf with a Dnmt1 antibody. Our results identified Dnmt1-positive EPCs migrating within the Dnmt1-positive population of endodermal cells and presumptive ISMPs that prefigure the intestinal epithelium (Fig. 3B). We conclude that EPCs and ISMPs express both *uhrf1* and *dnmt1*.

### Epigenetic modifiers Uhrf1 and Dnmt1 function together during intestinal tract development

The expression patterns of *dnmt1* and *uhrf1* showed that they are expressed in the same cells at the same time, suggesting the hypothesis that Uhrf1 and Dnmt1 function together during establishment of the intestinal tract. To test this hypothesis, we first examined the number of enteric neurons in *dnmt1^s904^* and *uhrf1* mutants at 5 dpf. The *dnmt1^s904^* mutation was isolated from an ENU mutagenesis screen for regulators of pancreas development. This allele is a splice acceptor mutation resulting in a complete loss of Dnmt1 catalytic activity (Anderson et al., 2009). We found that both *dnmt1* and *uhrf1* mutant larvae had similar reductions in ENS neurons (Fig. 4A,B). To further test interactions between Dnmt1 and Uhrf1, we crossed adult *dnmt1*;*uhrf1* double heterozygotes into a Tg(*phox2b*:*EGFP*) background and examined enteric neuron number. These double mutants displayed an ENS phenotype resembling that of *uhrf1* and *dnmt1* single mutants, and were not more severely affected (Fig. 4C). Our results from this epistasis analysis are consistent with the hypothesis that Uhrf1 and Dnmt1 act together during ENS development.

**Figure 4.**
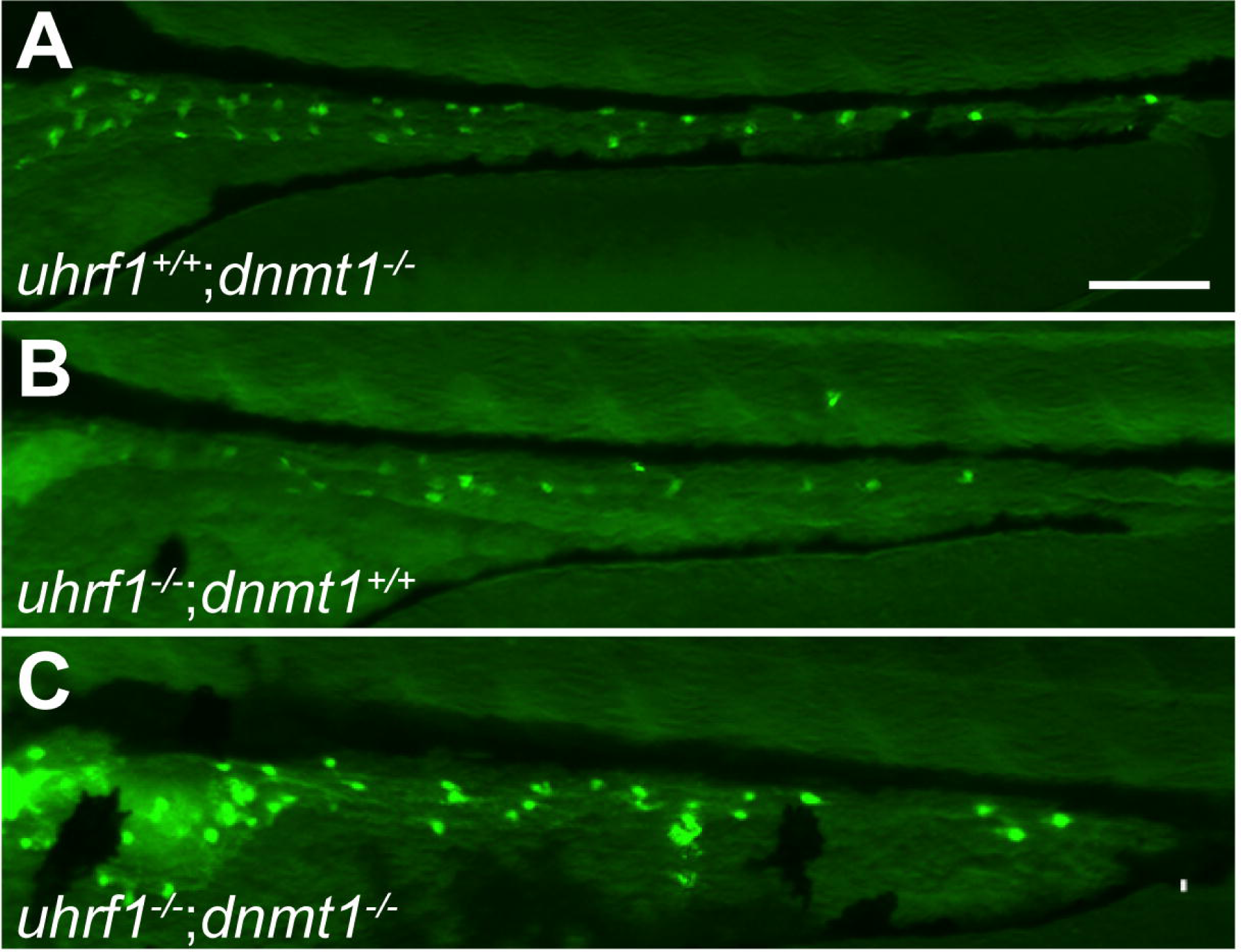
*uhrf1*;*dnmt1* double mutants show similar reduction in enteric neurons compared to *uhrf1* and *dnmt1* single mutants. (A) *dnmt1* single mutants show a similar reduction in *phox2b:EGFP* positive enteric neurons (green) compared to *uhrf1* single (B) or *uhrf1*;*dnmt1* double mutants (C). A-C: side views of whole-mount zebrafish larvae at 5 dpf. Scale bar = 50μm.

Homozygous *uhrf1* mutants were previously shown to have disrupted intestinal epithelium development (Marjoram et al., 2015). We found that *uhrf1* is expressed in presumptive ISMPs, prompting us next to test whether *uhrf1* mutants also show altered development of intestinal smooth muscle. We analyzed smooth muscle development in 5 dpf whole-mount *uhrf1* mutants and found that both longitudinal and circumferential intestinal smooth muscle fibers were essentially absent (Fig. 5A-D). Cross-sections stained with neuronal and smooth muscle markers or with Hematoxylin and Eosin (HE) revealed severe disruption of both intestinal smooth muscle and ENS components (Fig. 5E-L), and also showed that, instead of being sandwiched between the longitudinal and circumferential smooth muscle cell layers, enteric neurons of *uhrf1* mutants were often detached from the intestine (Fig. 5H). Interestingly, neuronal cell bodies, especially in the distal intestine of *uhrf1* mutants, had a rounded morphology compared to wildtype enteric neurons and also exhibited fewer processes, making them morphologically distinct from their wildtype counterparts (Fig. 5G, H).

**Figure 5.**
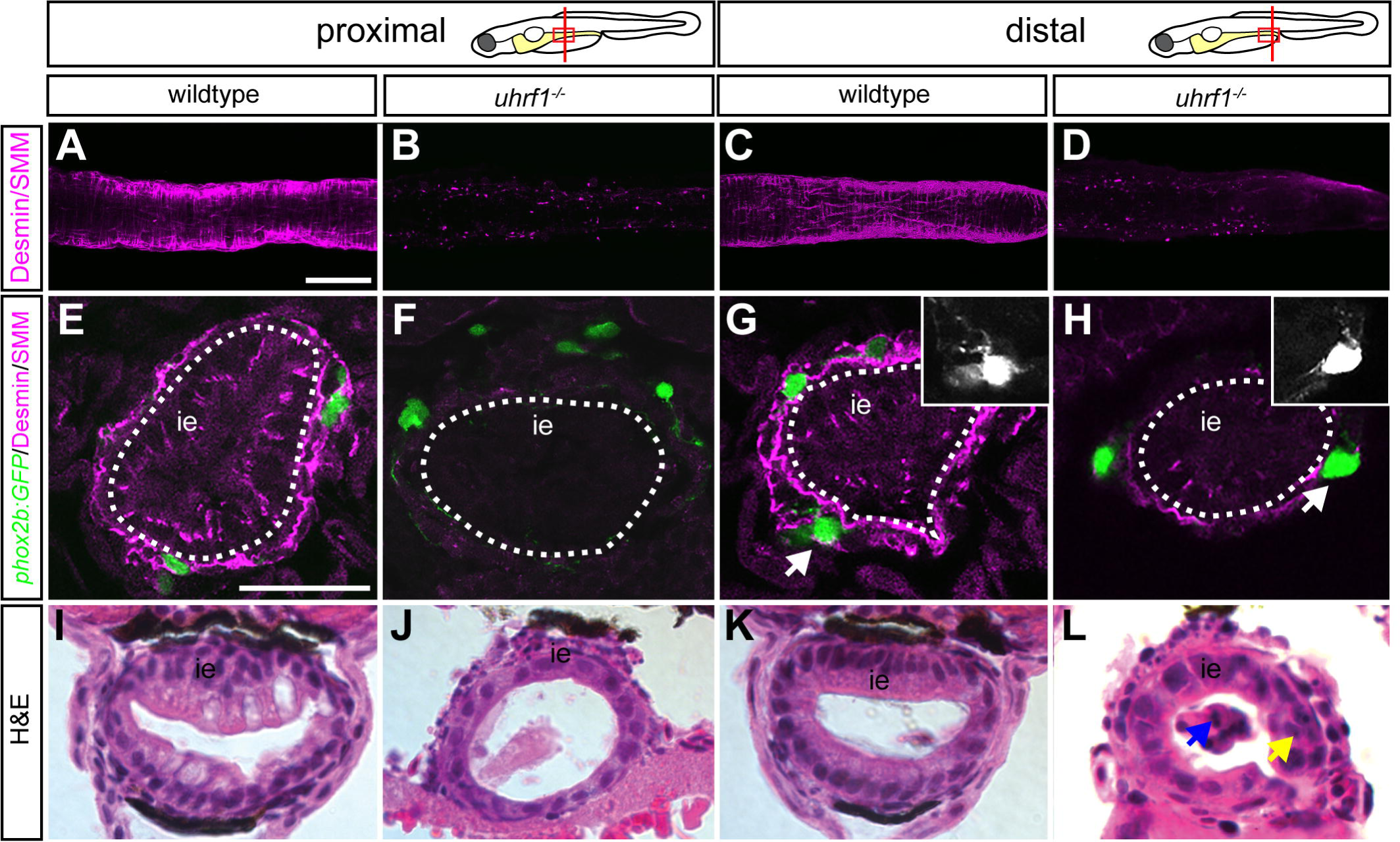
Smooth muscle cell and intestinal development is severely disrupted in *uhrf1* mutants. In both proximal and distal intestine, *uhrf1* mutants essentially lack smooth muscle cells, as revealed by labeling with smooth muscle myosin (SMM) and desmin (magenta) antibodies in whole-mounts (wildtype A,C *uhrf1^-/-^* B,D) and cross-sections (wildtype E,G, *uhrf1^-/-^* F,H). Close-ups in G and H indicate changes in neuronal morphology and displacement of *phox2b:EGFP*-positive neurons (green) in *uhrf1* mutants compared to wildtypes. Cross-sections stained with H&E of *uhrf1* mutants (J, L) show disrupted proximal (J) and distal (L) intestinal development compared to wildtypes (I, K). Increased shedding of cells in *uhrf1* mutants is indicated by a yellow arrow and disrupted intestinal epithelium with a blue arrow. ie = intestinal epithelium. A-D: confocal images of whole-mount dissected intestines; E-H: cross-sections at the levels indicated. I-L: brightfield images of cross-sections at the levels indicated. Scale bar = 50μm in A-D, and 25μm in E-L.

Dnmt1 has an essential role in smooth muscle cell development in mouse and humans (Jorgensen et al., 2018). To test whether this role of Dnmt1 is conserved in fish, we examined intestinal smooth muscle development in zebrafish *dnmt1* mutants. We found that, like *uhrf1* mutants, *dnmt1* mutants had severely disrupted intestinal smooth muscle development (Fig. 6). In addition, enteric neurons of *dnmt1* mutants shared the same rounded morphology as those of *uhrf1* mutants (Fig. 5). Together, the experiments described in this section show that Uhrf1 and Dnmt1 function together to control intestinal tract development by regulating development of each of the constituent cell types.

**Figure 6.**
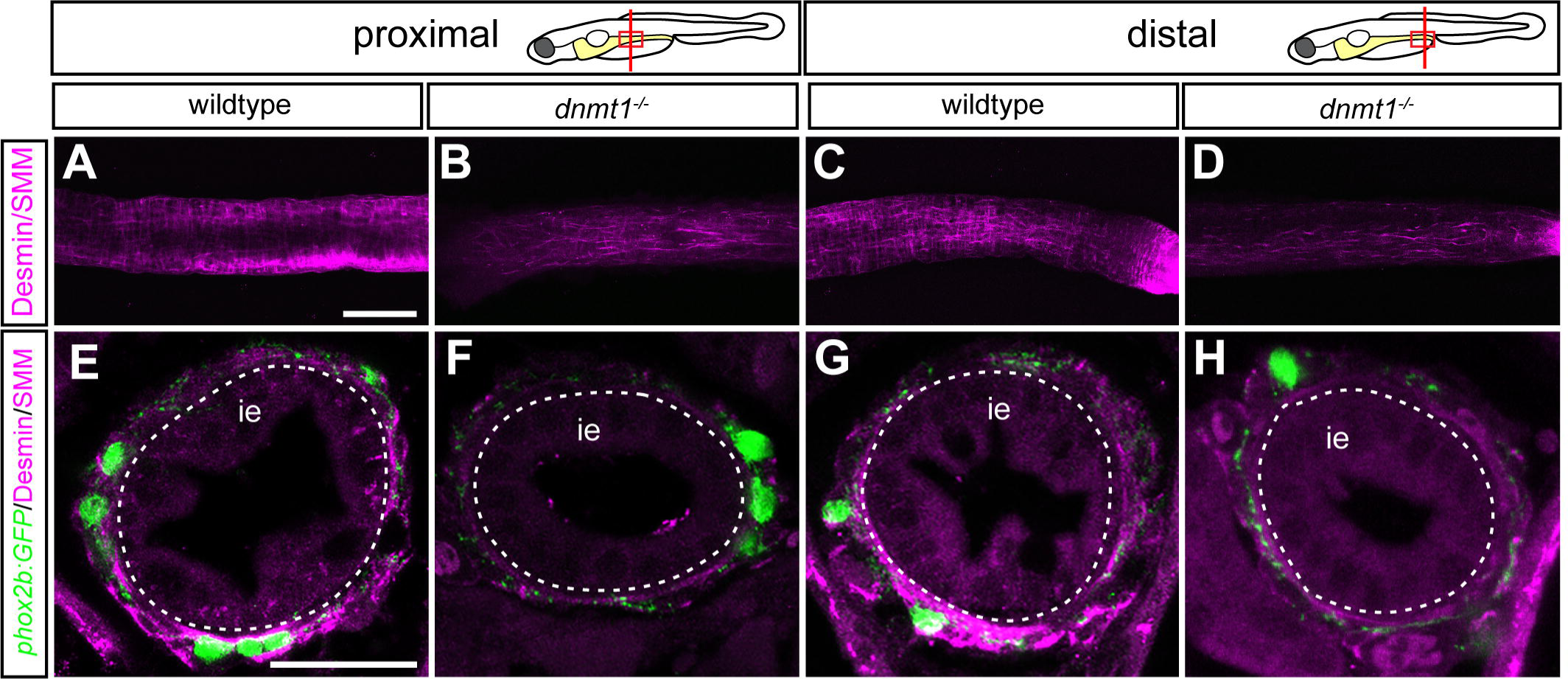
Smooth muscle cell and intestinal development is severely disrupted in *dnmt1* mutants. In both proximal and distal intestine, *dnmt1* mutants essentially lack smooth muscle cells, as revealed by labeling with smooth muscle myosin (SMM) and desmin (magenta) antibodies in whole-mounts (wildtype A,C *dnmt1^-/-^* B,D) and cross-sections (wildtype E,G, *dnmt1^-/-^* F,H). ie = intestinal epithelium. A-D: confocal images of whole-mount dissected intestines; E-H: cross-sections at the levels indicated. Scale bar = 50μm in A-D, and 25μm in E-L.

### Genetic chimeras reveal that *uhrf1* functions both cell-autonomously and cell-non-autonomously in ENS development

EPCs, smooth muscle precursors, and epithelial precursors are intermingled during their development and previous studies have shown that signals among these cell types are critical for proper differentiation of each cell type (Fu et al., 2004; Graham et al., 2017; Hao et al., 2016; Mwizerwa et al., 2011; Natarajan et al., 2002; Neunlist et al., 2007; Neunlist et al., 2013; Pietsch et al., 2006; Puzan et al., 2018; Reichenbach et al., 2008; Sukegawa et al., 2000). To learn whether loss of Uhrf1 disrupts interactions between cells derived from the different germ layers, we focused on EPCs. The disrupted development of both intestinal smooth muscle and intestinal epithelial cells was in *uhrf1* mutants, suggesting that Uhrf1 could act cell-non-autonomously during ENS development. To test this hypothesis, we generated genetic chimeras in which the vagal neural crest that contributes to the ENS was wildtype and the epithelium and muscle were mutant for *uhrf1* (Tables S1,S2). Chimeras were generated by transplanting vagal neural crest from *phox2b*:*EGFP*-expressing wildtype donors into host larvae from an incross of heterozygous *uhrf1* mutants. Thus, some hosts were wildtype, some were heterozygous, but phenotypically wildtype, and some were homozygous mutant (Fig. 7A). In this experimental set-up, all GFP-positive enteric neurons were donor-derived. As predicted, wildtype EPCs transplanted into phenotypically wildtype hosts contributed to the ENS along the entire length of the intestinal tract (Fig. 7B,C). In the context of a *uhrf1* mutant host environment, however, wildtype donor EPCs contributed to much of the intestinal tract, yet specifically were absent from the distal-most region of the intestine (Fig. 7C,D, asterisk, 5/5). This result suggests that Uhrf1 function is necessary cell-non-autonomously for complete EPC migration to form an ENS along the entire intestinal tract.

**Figure 7.**
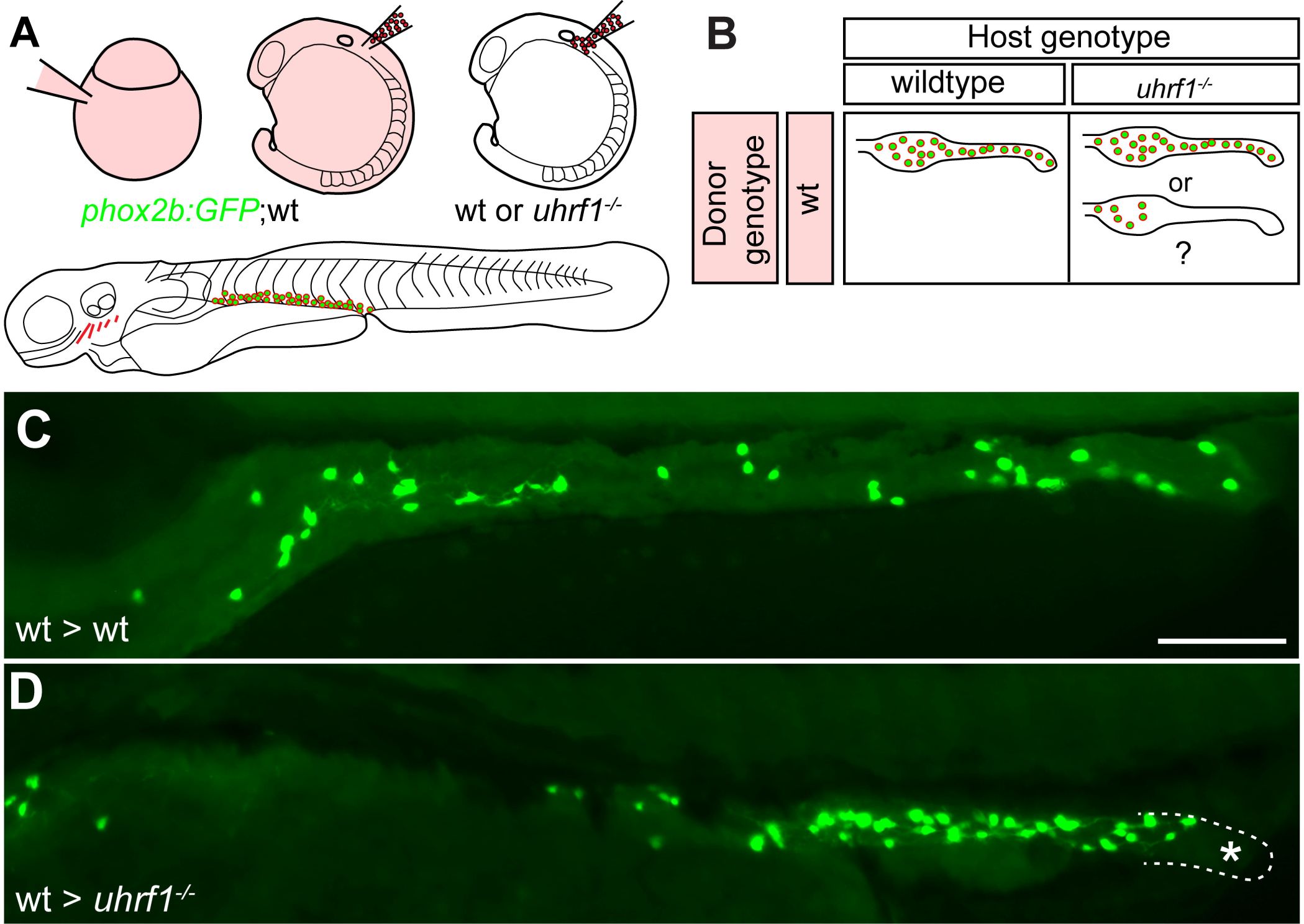
Genetic chimeras reveal that Uhrf1 functions cell-non-autonomously in ENS development. (A) Transplantation experiment diagram. Wildtype donor embryos that carry the *phox2b:EGFP* transgene were injected with rhodamine dextran. Vagal crest cells were transplanted from donors into unlabeled wildtype or *uhrf1* mutant hosts. The contribution of transplanted cells was evaluated at 5 dpf. (B) Possible results for transplantation experiments. (C) *phox2b:EGFP* positive (green) wildtype cells transplanted into wildtype hosts can populate the entire intestine. (D) *phox2b:EGFP* positive (green) wildtype cells transplanted into *uhrf1* mutant hosts can expand, but are excluded from distal intestine (outlined and labeled with asterisk). C, D: lateral views of whole-mount zebrafish larvae at 5 dpf. Scale bar = 50μm in C,D.

Because *uhrf1* is expressed in EPCs, we also tested whether Uhrf1 function is required cell-autonomously. We performed the same transplantation as described above, but transplanted vagal neural crest from labeled donor embryos derived from a *uhrf1* mutant incross carrying the *phox2b*:*EGFP* transgene into unlabeled wildtype hosts (Fig. 8A, Table S3). As in the previous experiment, wildtype donor cells transplanted into wildtype hosts populated the entire length of the intestine (Fig. 8B,C). In contrast, in most cases mutant donor EPCs migrated only a short distance caudally after entering the wildtype host intestine (4/5). Even in the single case in which transplanted *uhrf1* mutant cells reached the distal intestine (1/5), the mutant donor cells displayed mutant morphology and the precursors did not proliferate as much as wildtype precursors (Fig. 8D-F), showing that Uhrf1 functions cell-autonomously in EPCs.

**Figure 8.**
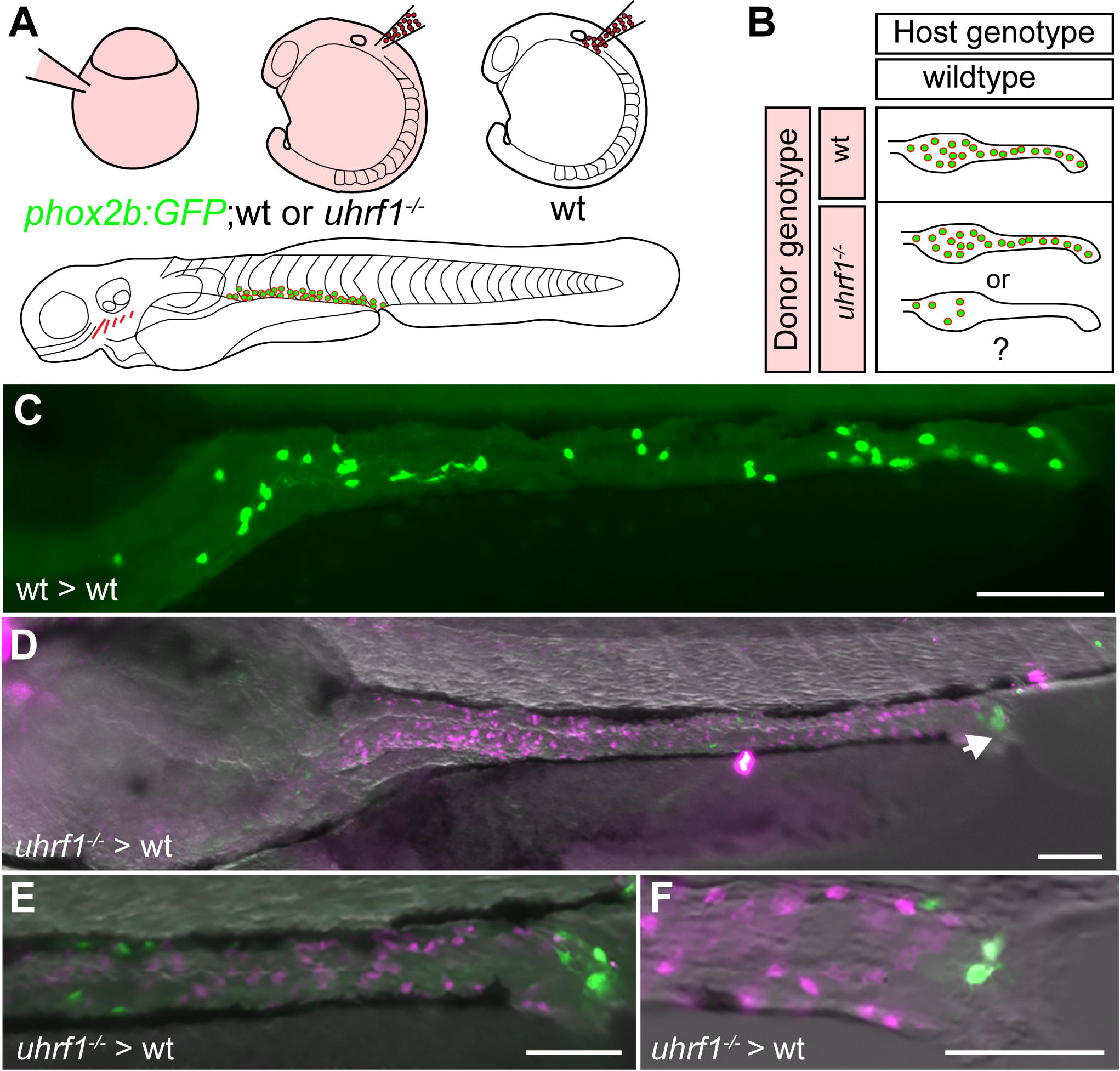
Genetic chimeras reveal that Uhrf1 functions cell-autonomously in ENS development. (A) Diagram of transplantation experiment. Wildtype or *uhrf1* mutant donors that carry the *phox2b:EGFP* transgene were injected with rhodamine dextran. Vagal crest cells were transplanted from donors into unlabeled wildtype hosts. The contribution of transplanted cells was evaluated at 5 dpf. (B) Possible results for transplantation experiments. (C) *phox2b:EGFP* positive (green) wildtype cells transplanted into wildtype hosts can populate the entire intestine. Please note that we included the same wt > wt images as in Fig. 7 for clarity. (D) *phox2b:EGFP* positive (green) *uhrf1* mutant cells transplanted into wildtype hosts can migrate to distal intestine (1/5), but they do not expand as much as wildtype cells. (E) close-up of posterior region of the gut shown in (D). (F) *phox2b*:EGFP positive (green) *uhrf1* mutant cells (arrows) only migrate to the anteriormost portion of the intestine (4/5) and they do not expand. Intestine outline with dashed lines in F. Note that Elavl positive cells (magenta) that are EGFP negative are host-derived enteric neurons. C-F: lateral views of whole-mount zebrafish larvae at 5 dpf. Scale bar = 50μm in C-F.

## Discussion

This study provides evidence that disruption of DNA methylation factors interrupts the developmental trajectories of the three distinct cell types that form the intestinal tract, thus preventing formation of a functional organ. Previous studies have demonstrated that these three cell types, intestinal epithelium, intestinal smooth muscle, and the cells of the enteric nervous system, develop over a similar time frame. Importantly, cells derived from these distinct germ layers provide signals that are critical for one another’s differentiation (Fu et al., 2004; Graham et al., 2017; Hao et al., 2016; Mwizerwa et al., 2011; Natarajan et al., 2002; Neunlist et al., 2007; Neunlist et al., 2013; Pietsch et al., 2006; Puzan et al., 2018; Reichenbach et al., 2008; Sukegawa et al., 2000) (Fig. 9). Mouse *Uhrf1* mutants die early, before the intestine develops (Bostick et al., 2007; Muto et al., 2002; Sharif et al., 2007), and so mouse does not serve as a model in which to investigate the role of Uhrf1 in intestinal development. In contrast, zebrafish *uhrf1* mutants survive to larval stages due to maternally provided Uhrf1 protein (Jacob et al., 2015), thus the role of *uhrf1* in intestinal development can be investigated._Below we discuss the role of methylation factors in the context of intestinal development and disease.

**Figure 9.**
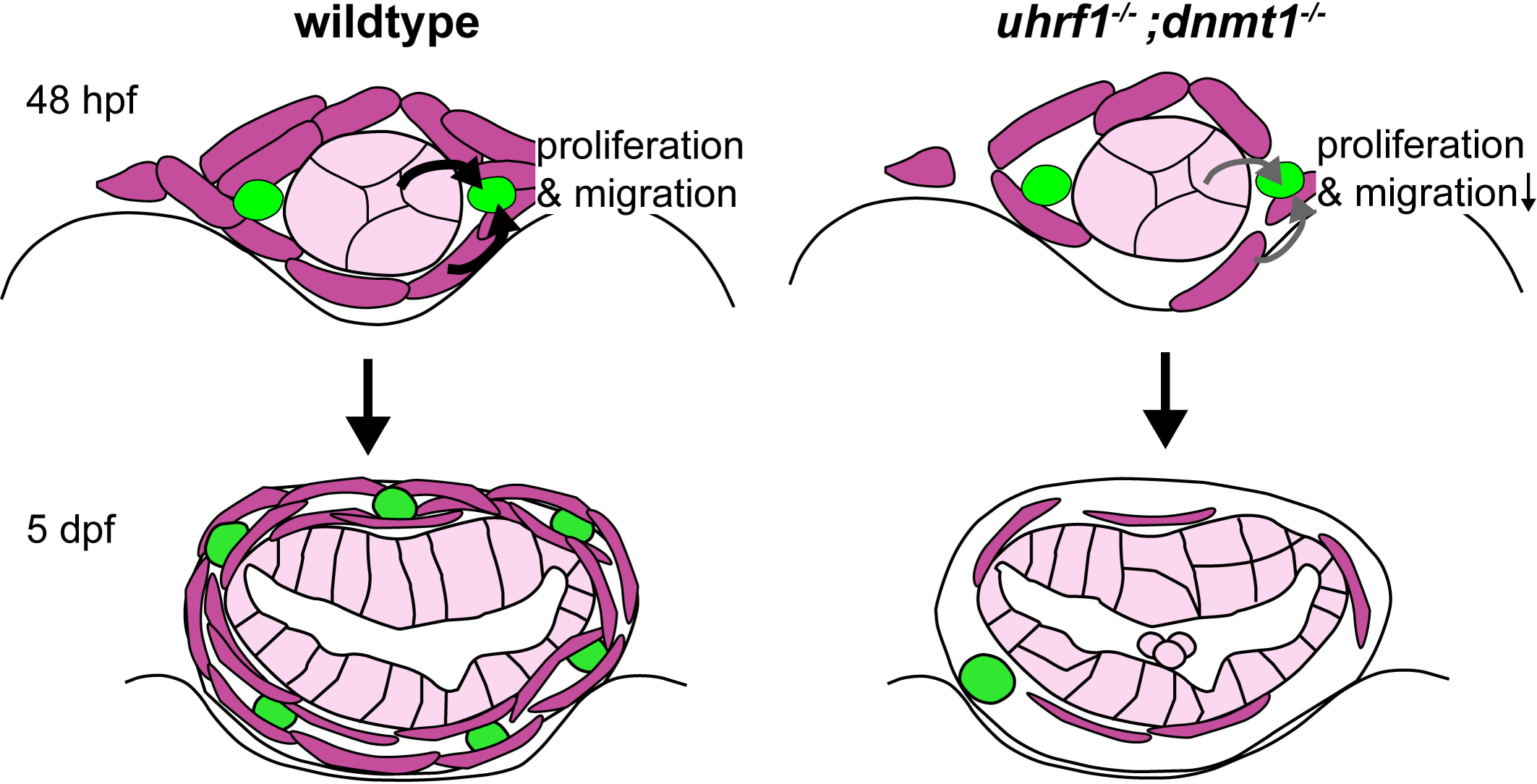
Model of Uhrf1 and Dnmt1 function during intestinal development. During early intestinal development, Uhrf1 and Dnmt1 function cell-autonomously in EPCs (green) and cell-non-autonomously in surrounding intestinal epithelial cells (light pink) and smooth muscle cells (purple). Signaling is important for differentiation of cells derived from each of the germ layers. This diagram shows signaling from epithelial and muscle precursors to ENS precursors. Decreasing these signals in *uhrf1* and *dnmt1* mutants results in fewer smooth muscle and epithelial progenitors, and thus alters proliferation, migration, or differentiation of EPCs.

### Uhrf1 and Dnmt1 promote coordinated intestinal development

Development of a functional intestine is the result of highly orchestrated developmental processes. Each of the contributing germ layers must be specified and component cells must segregate and migrate to their appropriate destinations (Ganz, 2018; Ganz et al., 2016; Hao et al., 2016; Wallace et al., 2005a). Endoderm reaches the midline early and forms a scaffold for the incoming mesoderm that migrates from the lateral plate (Gays et al., 2017). In contrast, the ectoderm-derived neural crest cells first migrate ventrally to the developing intestine and then migrate from rostral to caudal along the developing intestinal epithelium and mesoderm. All of these processes occur within a fairly narrow time window. During migration, each cell type also proliferates, a process that has been best studied for the ENS (Shepherd and Eisen, 2011). In the ENS, migration and proliferation are intimately connected; decreased proliferation leads to decreased migration and vice versa (Landman et al., 2007; Nagy and Goldstein, 2017; Newgreen et al., 2017; Simpson et al., 2007). Uhrf1 and Dnmt1 mutations block the cell cycle, leading to fewer cells in the affected tissue or organ (Jacob et al., 2015), providing a mechanism for the decreased number of ENS neurons in these mutants. A similar mechanism may be at play in intestinal smooth muscle and intestinal epithelium, although this hypothesis remains to be tested. Because each cell type independently fails to form properly, signals produced by that cell type that are necessary for differentiation of the other two cell types decrease. We cannot, however, entirely rule out the possibility that some EPCs, ISMPs, or intestinal epithelial progenitor cells differentiate prematurely and thus fail to migrate properly. However, even in this case, because there would be fewer cells and they would not assume their normal positions, inter-germ layer signaling would likely be decreased. Mutations in genes involved in methylation can also keep progenitor cells in an undifferentiated, proliferative state (Sen et al., 2010), another possible mechanism for decreasing the number of cells that would lead to a similar decrease in proper migration and thus a diminution of signals important for adjacent cell types to differentiate.

Our transplantation experiments provide evidence that methylation factors act both cell-autonomously and cell-non-autonomously in the ENS and we suggest that this is also likely to be the case in the intestinal smooth muscle and epithelium. This hypothesis can be tested by creating genetic chimeras for these cell populations or by cell type specific knock outs.

### Uhrf1 and Dnmt1 support healthy intestinal development

Epigenetic regulation of gene expression has been implicated in diseases that affect the intestinal epithelium, intestinal smooth muscle, and ENS. For example, dysregulation of the intestinal epithelium has been implicated in inflammatory bowel disease (IBD) as a result of hypoproliferation and excess shedding of epithelial cells into the intestinal lumen (Marjoram et al., 2015). However, in the study by Marjoram and colleagues (2015), *uhrf1* or *dnmt1* were mutant throughout the organism, including in all cells of the intestinal tract, but contributions from either intestinal smooth muscle or ENS progenitors to the reported phenotype were not examined. The ENS has been implicated in regulating intestinal barrier function (Neunlist et al., 2013; Sharkey, 2015), suggesting that the ENS phenotype observed in our study could be a contributing factor to compromised intestinal barrier function in *uhrf1* mutants. This explanation may also help elucidate why not all aspects of the intestinal phenotype were restored in rescue experiments (Marjoram et al., 2015). *Dnmt1* mutations engineered to be present solely in mouse intestinal smooth muscle have been shown to result in muscle hypoproliferation and decreased intestinal peristalsis, phenotypes found in human diseases such as chronic intestinal pseudo-obstruction and megacystis-megacolon-intestinal hypoperistalsis syndrome (Jorgensen et al., 2018). Interestingly, DNMT1 is dysregulated in one type of IBD, Crohn’s disease (Jorgensen et al., 2018). DNA hypomethylation has also been implicated in ENS-mediated diseases, particularly HSCR and Waardenburg syndrome (Kim et al., 2011; Torroglosa et al., 2016; Torroglosa et al., 2014). Changes in maintenance of DNA methylation can change gene expression patterns and several HSCR loci including *RET, EDNRB*, and HSCR candidate genes, such as *PAX6*, have been shown to be affected by changes in DNA methylation in their promoters (Torroglosa et al., 2016). Epigenetic modulation of methylation is also linked to other intestinal diseases, for example intestinal cancers (Gays et al., 2017). Zebrafish *uhrf1* mutant alleles have been demonstrated to decrease DNA methylation (Marjoram et al., 2015; Tittle et al., 2011). Thus, our results provide strong evidence that DNA methylation is critical for normal development of each of the cell types that constitute the intestine, and that failure in the methylation process can lead to devastating disruptions of cell type development that manifest in a variety of disease states to which these cell types contribute.

## Materials and Methods

### Zebrafish husbandry

All zebrafish experiments were performed following protocols approved by the University of Oregon Institutional Animal Care and Use Committee. Homozygous *uhrf1^b1115^* and *dnmt1^s904^* single mutants were obtained by mating heterozygous carriers. Complementation testing was performed by mating adult zebrafish heterozygous for *uhrf1^b1115^* and *uhrf1^hi272^* (Sadler et al., 2007). Mutant larvae were first identified visually by smaller eyes and jaws and subsequently confirmed by fixing and staining with the pan-neuronal marker anti-Elavl and analyzing neuronal numbers in the intestine. For some experiments, *uhrf1^b1115^* and *dnmt1^s904^* mutant carriers were outcrossed to *Tg(phox2b:EGFP)^w37^* (Nechiporuk et al., 2007) and their progeny were grown up and screened for single and double mutant carriers that were subsequently crossed to visualize EPCs and neurons in living animals.

### RAD-tag genotyping

Previously we performed a forward-genetic screen to uncover regulators of neural crest development (Kuhlman and Eisen, 2007). One of the mutants identified, *b1115,* displayed a severe enteric neuron phenotype [Fig. 1, (Kuhlman and Eisen, 2007)]. Using Restriction site Associated DNA (RAD) sequencing of genomic DNA (Baird et al., 2008), we mapped the *b1115* genetic lesion to a 2.2 Mb region on chromosome 22. Phenotypic mutant and wildtype fish from a single-pair cross were identified by *phox2b:EGFP* transgene expression in the ENS at 5 dpf and 5 mutant or 5 wildtype individuals were pooled for sequencing. Genomic DNA was purified from the cross parents, from 9 wildtype larval pools, and from 34 *b1115* mutant larval pools. RAD-tag libraries were prepared as described (Amores et al., 2011; Baird et al., 2008; Miller et al., 2007) and sequenced on an Illumina HiSeq 2000 to obtain 100-nucleotide single-end reads. We used Stacks version 1.19 (http://catchenlab.life.illinois.edu/stacks/) to organize reads into loci and to identify polymorphisms (Catchen et al., 2013; Catchen et al., 2011). Illumina sequences were quality filtered with the process_radtags program of Stacks (Catchen et al., 2013; Catchen et al., 2011) and aligned to the zebrafish genome (v. Zv9) (Howe et al., 2013) using GSNAP (Wu and Nacu, 2010). We ran pstacks with default parameters on the parent sequences, and with the parameters --bound_high.001 and --alpha. 001 on pooled samples. The Stacks catalog was built from the parent sequences and sample stacks were matched to the catalog with sstacks. We excluded all loci with indels and more than two alleles in a single locus, and used the program SNPstats to calculate a G-test statistic comparing genotypes in mutants and wildtype pools (Hohenlohe et al., 2010). Sequences have been deposited in the Sequence Read Archive (https://www.ncbi.nlm.nih.gov/sra) under the accession number SRP118720.

Analysis of genes within the interval implicated *uhrf1* as a candidate due to the similarities in phenotype between *uhrf1^b1115^* and the previously described transgene insertion allele *uhrf1^hi272^* (Amsterdam and Hopkins, 2004; Sadler et al., 2007). Crossing the *b1115* allele to the *hi272* allele resulted in non-complementation (Fig. S1E,F) indicating that *b1115* is a mutated allele of *uhrf1*. To learn the site of the genetic lesion, we generated cDNA from pools of wildtype and mutant larvae, amplified and sequenced the *uhrf1* gene in each pool and performed sequence comparisons. We identified a single base pair change resulting in a cysteine-to-glycine change in amino acid 744, which lies in the zinc finger RING domain of the protein (Fig. S1A-C). Sequence analysis of vertebrate species in the conservation track of the human-centric 100-way species alignment in the UCSC Genome Browser (https://genome.ucsc.edu/; representative species shown in Fig. S1D) revealed that the affected amino acid is invariably conserved in all vertebrates in the alignment and that the affected cysteine (C7) is an essential part of the RING finger domain involved in binding of one of the zinc ions that are critical for the function of the RING finger domain (Borden and Freemont, 1996) (Fig. S1D).

### Immunohistochemistry

Antibody staining for Elavl (1:10,000, Thermo Fisher Scientific, Eugene, OR, catalog number A-21271), GFP (1:1000, Thermo Fisher Scientific, Eugene, OR, A11122, catalog number A11120, nNOS (1:500, Santa Cruz Biotechnology, Santa Cruz, CA, catalog number sc-1025), 5HT (1:10,000, Immunostar, Hudson, WI, catalog number 20080), Smooth muscle myosin (1:100, Alfa Aesar, catalog number J64817 BT-562), and Desmin (1:100, Sigma-Aldrich, catalog number D8281) was performed at 5 dpf as previously described (Uyttebroek et al., 2010). Antigens were visualized with standard fluorophore-labeled antibodies for rabbit IgG (1:1,000, Thermo Fisher Scientific, Eugene, OR, catalog number A-11008 or A-11071) and mouse IgG (1:1,000, Thermo Fisher Scientific., Eugene, OR, catalog number A-11001 or A-11030).

### Hematoxylin and Eosin staining

Embryos were fixed in 4% paraformaldehyde and processed for paraffin sectioning as previously described (Cheesman et al., 2011).

### *In situ* hybridization

Embryos were fixed and processed for *in situ* hybridization as previously described (Seredick et al., 2012). Following *in situ* hybridization, embryos were embedded in agar, frozen, cryosectioned (Beattie and Eisen, 1997), and imaged on a confocal microscope as described below.

### ENS Transplantation

ENS transplantation was performed as previously described (Rolig et al., 2017). Briefly, wildtype donor embryos were labeled by injection of 5% tetramethylrhodamine dextran (3000 MW) at the 1-2 cell stage and reared until the next manipulation in filter-sterilized embryo medium (EM). Embryos at the 12-14 somite stage were mounted in agar, a small hole dissected in the skin, and cells transplanted as previously described (Eisen, 1991). At 5 dpf, hosts and donors were fixed in 4% paraformaldehyde in 1× PBS for 3 h at room temperature and then washed 3 × 10 min in 1× PBS. We performed immunohistochemistry with anti-Elavl and anti-GFP (see immunohistochemistry section) to reveal both host and donor-derived enteric neurons.

### Image acquisition

Confocal images were acquired on a Zeiss Pascal confocal microscope using a 40X water immersion objective and AIM or ZEN software (Carl Zeiss Microscopy, LLC, Thornwood, New York, USA) or a Leica SP8 confocal microscope using a 40X water immersion objective or a 63X oil immersion objective, 405-diode, white light lasers and LASX software. Low magnification images were acquired using a Leica MZ205 FA fluorescent stereomicroscope and LAS software. Images were processed and analyzed using Photoshop CC (Version 19.1.1, Adobe Systems, Inc., San Jose, CA, USA) and FiJi software.

### Imaging analysis

Following immunohistochemistry, intestines were dissected and mounted in PBS between two coverslips. For cell quantification, projections along the y-axis were generated from confocal stacks in ZEISS LSM Image browser and used for manual cell counts and colocalization analysis between Elavl and nNOS or Elavl and 5-HT. Counts were performed in proximal intestine, defined as extending 200 μM from the caudal end of the intestinal bulb, and in the distal intestine, defined as starting 200 μM from the anus.

## Acknowledgments

We thank Doug Turnbull and the University of Oregon Genomics and Cell Characterization Core Facility for their expertise in Illumina sequencing and bioinformatics, Poh Kheng Loi for histology, Adam Christensen, John Dowd and the UO Zebrafish Facility staff for assistance with zebrafish husbandry.

## Author contributions

Conceptualization: JSE, JG, EM

Methodology: JSE, JG, EM, AA, CW, JHP

Validation: JG, EM, AA, CW, JHP, MS, PB, IB

Formal analysis: JG, EM, AA, CW, JHP, MS, PB, IB, JSE

Investigation: JG, EM, AA, CW, PB, MS, IB, PD, JK, JHP, JSE

Resources: JSE, JHP, JG

Data curation: CW, EM, JG, JSE

Writing-original draft preparation: JG, EM, JSE

Writing-review and editing: JG, EM, AA, CW, PB, MS, IB, PD, JK, JHP, JSE

Visualization: EM, JG, JSE

Project administration: JSE

Funding acquisition: JSE, JHP, JG

Supervision: JSE, JHP

Mutant mapping: EM, JG, AA, CW, PB, IB

---

**Figure S1. The zebrafish *b1115* mutation has a non-synonymous single nucleotide polymorphism in an ultra-conserved amino acid of the RING domain of *uhrf1*.** (A) Exon 16 of *uhrf1* contains the mutation (red asterisk) in the *b1115* allele. (B) Schematic of important protein domains of *uhrf1* with a close-up of the 7 conserved Cysteines. (C) A non-synonymous single nucleotide polymorphism (T=>C) mutates the conserved Cysteine 7 (red) to a Glycine. (D) the cysteine 7 (red) is conserved among phylogenetically diverse vertebrate species.(E) Complementation crosses between the *b1115* allele and the *hi272* allele of uhrf1 show that the two alleles fail to complement as they have the mutant phenotype of fewer *phox2b*:EGFP enteric neurons (F, green) compared to wildtype (E). E,F: side-views of whole-mount zebrafish larvae at 5 dpf. Scale bar = 100μm in E,F.

**Table S1.** Transplantation analysis of wildtype *phox2b:EGFP* vagal neural crest cells to *uhrf1^-/-^* hosts at 5 dpf. Transplanted cell morphology, cell position and relative numbers were recorded. The letter y denotes mutant cellular morphology, y=1-5 cells, yy=6-20 cells, yyy= 20 or more cells. Each row represents an individual larva.

| Transplanted cell location |  |  |  |
| --- | --- | --- | --- |
| Esophageal Intestinal Junction | Bulb | Proximal intestine | Distal intestine |
|  | y | yyy |  |
|  | y | yyy |  |
|  | y | yyy |  |
|  | y | yy |  |

**Table S2.** Transplantation analysis of wildtype *phox2b:EGFP* vagal neural crest cells to wildtype hosts at 5dpf. Transplanted cell morphology, cell position and relative numbers were recorded. The letter x denotes wildtype cellular morphology, x=1-5 cells, xx=6-20 cells, xxx= 20 or more cells. Each row represents an individual larva.

| Transplanted cell location |  |  |  |
| --- | --- | --- | --- |
| Esophageal Intestinal Junction | Bulb | Proximal intestine | Distal intestine |
|  |  | x | x |
|  |  | x |  |
|  | xx | xx |  |
|  | xxx | xxx | xxx |
|  | xxx | xx | xx |
|  |  | xx | xx |
|  | x | xxx | x |
|  | xx | xx | xx |
|  | xx | xx | xx |
|  | x | x | x |
|  |  | xx | xx |
|  | x | x | x |
|  | xxx | xxx | xxx |
|  |  |  | xx |
|  |  | xx | xxx |
| x | xx |  |  |
|  | xx | xx | xx |
| x | xx | xx | xx |
|  | xx | xx | xx |

**Table S3.** Transplantation analysis of *uhrf1^-/-^; phox2b:EGFP* vagal neural crest cells to wildtype hosts at 5dpf. Transplanted cell morphology, cell position and relative numbers were recorded. The letter y denotes mutant cellular morphology, y=1-5 cells, yy=6-20 cells, yyy= 20 or more cells. Each row represents an individual larva.

| Transplanted cell location |  |  |  |
| --- | --- | --- | --- |
| Esophageal Intestinal Junction | Bulb | Proximal intestine | Distal intestine |
|  | y | y | y |
|  | y |  |  |
|  | y |  |  |
| yy |  | y |  |

